# Single-molecule imaging of DNA repair and cytoplasmic rigidification in intracellular bacteria

**DOI:** 10.64898/2026.09.14.751380

**Authors:** Fiona Anne Sargison, Nicolas Chérif Ducrot, Clara Kummerer, Claire Qu, Treasa B O’Hagan, Elliott Haywood, Anju Kudhail, Amy Moores, Lindsay Baker, Stephan Uphoff

## Abstract

DNA damage is an important component of the antibacterial response of phagocytes, but which DNA repair mechanisms are active in intracellular bacteria remains unclear. We developed a live-cell single-molecule tracking approach to directly measure DNA repair activity in *Escherichia coli* within macrophages. Phagocytosis activates bacterial base excision and nucleotide excision repair pathways and increases DNA mismatch repair foci indicative of DNA replication errors. Phagocyte-generated stresses also cause a general slowdown in protein diffusion within bacteria, consistent with a transition of the cytoplasm towards a glass-like state, which is associated with reduced metabolic activity. At the single-cell level, DNA repair activity is highly heterogeneous, with metabolically inactive bacteria showing the greatest engagement of repair proteins. Together, these findings reveal how distinct DNA repair pathways are deployed during macrophage infection and link repair activity to the metabolic and biophysical states of individual intracellular bacteria.

## Introduction

Bacteria must withstand a range of stresses to survive within a host, particularly upon encountering the host’s immune defences. Professional phagocytes act at the frontline of the innate immune response, ingesting bacteria and exposing them to a combination of antimicrobial insults. While many bacteria are rapidly eliminated following phagocytosis, those that persist must cope with starvation and extensive host-generated damage^1,2^. Bacterial species classified as intracellular pathogens have evolved specialised strategies to acquire nutrients, and to remodel and detoxify the intracellular environment, enabling long-term survival and replication within phagocytes^1^.

However, before such adaptive mechanisms are established, bacteria first rely on basal defence systems that are broadly conserved across pathogenic and commensal species. Understanding this early phase of host-pathogen interaction is therefore critical, as it defines the fundamental protective mechanisms to withstand host-imposed stresses. Central to this basal bacterial defence is the ability to repair DNA damage^3^. After phagocytosis, bacteria are trapped in a progressively toxic, acidic, and nutrient-deprived phagosome, where limited energy availability may constrain detoxification and repair processes^2^. In parallel, the phagocyte oxidative burst exposes bacteria to reactive oxygen and nitrogen species (ROS, RNS) that can generate a spectrum of DNA lesions, including oxidised bases, abasic sites, and helix-distorting damage. Such lesions are known to be substrates for conserved DNA repair pathways, including base excision repair (BER), nucleotide excision repair (NER), mismatch repair (MMR), and double-strand break repair (DSBR) pathways^4,5^. Genetic studies have shown that disruption of these pathways compromises bacterial survival and genome stability during residence in phagocytes, and attenuates virulence in diverse pathogens such as *Escherichia coli*, *Salmonella enterica*, *Staphylococcus aureus,* and *Mycobacterium tuberculosis*^6–10^.

The stresses imposed by immune defences do not affect all bacteria uniformly. Even genetically identical cells adopt diverse phenotypic states in response to stress^11^. This diversification is further amplified by variability in the host cell environment and feedback between bacterial and immune responses, leading to divergent fates ranging from active growth to dormancy or death. Such heterogeneity poses a major challenge for bacterial clearance by immune defences. *In vitro*, single-cell studies have shown that DNA damage responses can themselves be highly heterogeneous, and analyses of intracellular bacteria have revealed cell-to-cell variation in the expression of DNA repair genes^12–15^. However, differences in protein abundance do not necessarily reflect differences in molecular activity or functional outcomes. Most insights into bacterial DNA repair during host infection derive from indirect measurements, typically based on the survival of mutant strains or endpoint analyses of bacteria extracted from host cells^16–18^. While such approaches establish the importance of specific genes, they provide limited information about when and where DNA repair occurs within intact host cells and how repair activity relates to the physiological state of individual bacteria and their local environment. It therefore remains unclear whether intracellular bacteria exhibit heterogeneous DNA repair activity, and how such variation relates to bacterial fate during infection. Addressing these questions requires approaches capable of resolving molecular activities in individual bacteria within host cells.

Here, we establish a live-cell imaging approach to directly visualise protein activity in bacteria residing within host cells. Single-molecule tracking (SMT) enables quantification of protein localisation, diffusion behaviour, and interactions *in situ*^19^. Using non-pathogenic *E. coli* as a model, we show that BER, NER, and MMR pathways are engaged after phagocytosis by macrophages and function to repair mutagenic DNA damage. Unexpectedly, SMT also showed that host-generated stresses drive a pronounced general slowdown in protein diffusion, indicating a transition of the bacterial cytoplasm toward a glass-like state associated with reduced metabolic activity. Single-cell analysis revealed that DNA repair activity varies with metabolic state, as inactive bacteria show higher engagement of repair proteins. Together, this work reveals how bacteria deploy DNA repair to defend against phagocyte attack and establishes a general method for probing molecular processes in bacteria during intracellular infection.

## Results

### Development of a method for single-molecule tracking of proteins within intracellular bacteria

SMT is an established method for measuring DNA repair processes in bacteria exposed to stress treatments under *in vitro* conditions that mimic aspects of a host cell environment^14,20^. The method reveals how repair proteins search for lesions and becomes transiently immobilised while they perform repair. Such reductionist systems, however, do not capture the complexity of bacteria-host interactions. Extending single-molecule imaging from controlled *in vitro* systems to bacteria within host cells is a natural next step but presents substantial technical challenges: cellular autofluorescence and light scattering reduce signal-to-noise, and intracellular bacteria are sparse and unevenly distributed. To overcome these limitations, we leveraged the genetically encoded HaloTag as a strategy for labelling specific proteins in live intracellular bacteria with bright and photostable fluorophores^21,22^. We used human macrophages differentiated from THP-1 monocytes as a standard infection model and *E. coli* MG1655, a well-characterised non-pathogenic laboratory strain in which conserved DNA repair pathways were discovered and extensively studied. We fused the native chromosomal gene of DNA polymerase I to HaloTag (Pol1-Halo). Pol1 performs gap-filling DNA synthesis during excision repair, making it a functional reporter of DNA lesion processing^3^. Pol1-Halo was labelled with TMR dye, excess dye was removed, and macrophages were infected at a multiplicity of infection (MOI) of 10 bacteria per cell (Fig 1a). Movies of intracellular bacteria recorded with an exposure time of 15 ms per frame. Using a highly inclined laser illumination, the real-time motion of individual Pol1-Halo-TMR molecules was captured.

**Figure 1.**
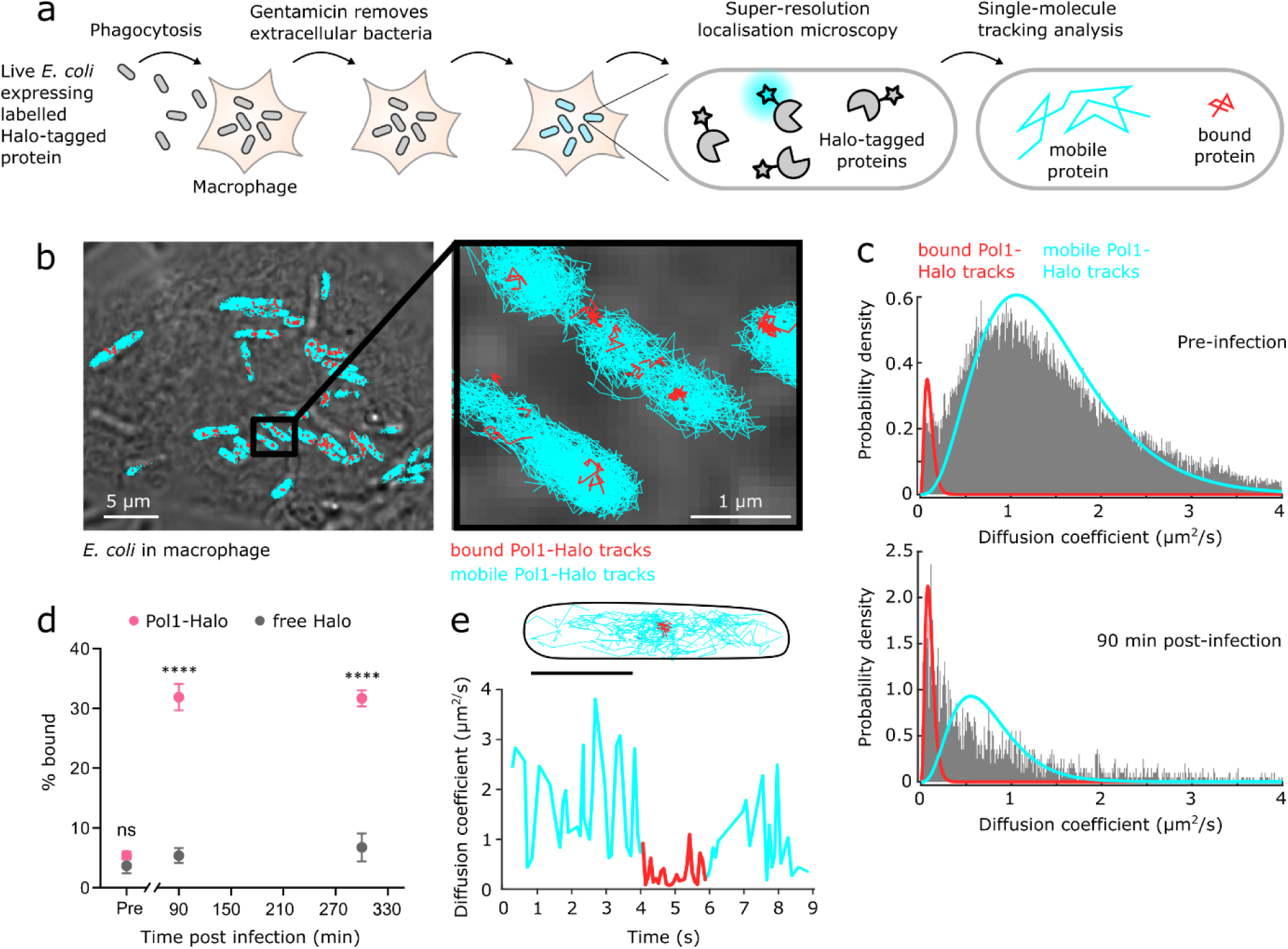
Single-molecule tracking of DNA repair polymerase Pol1 in *E. coli* within macrophages. **(a)** Schematic of the SMT workflow used to investigate DNA repair protein dynamics in intracellular bacteria. **(b)** Representative transmitted light image of a macrophage with overlay of Pol1-Halo tracks in intracellular *E. coli.* Tracks of Pol1 molecules classified as mobile or immobile shown in turquoise or red, respectively. **(c)** Histogram distributions of diffusion coefficients for Pol1-Halo molecules in *E. coli* pre-infection and at 90 min post-infection. A two-state diffusion model was fitted to estimate the relative fractions of immobile (red) and mobile (turquoise) molecules. Combined data from 3 independent repeats. **(d)** Percentages of immobile Pol1-Halo and free Halo molecules in *E. coli* pre-infection and at indicated times post-infection. Error bars indicate mean ± S.E.M. from 3 independent repeats. Comparison between Pol1-Halo and free Halo using two-way ANOVA with Dunnett’s multiple-comparisons test at each time point; ns, not significant; \*\*\*\**P* < 0.0001. **(e)** Representative time-trace of diffusion coefficient and corresponding track of single Pol1-Halo-JFX650 molecule showing transition between mobile (turquoise) and immobile (red) states. Scale bar = 1 µm

We observed a mixture of mobile and immobile protein tracks (Fig 1b, c). For DNA repair enzymes, immobile molecules typically reflect engagement at sites of DNA damage^14,19,20^. We found that the fraction of immobile Pol1 molecules increased markedly following phagocytosis, rising from an average of 5.65% in extracellular bacteria to 32.4% by 90 minutes after phagocytosis and remaining elevated at 31.7% at 300 min post-phagocytosis (Fig. 1c, d). These data indicate sustained excision repair activity during intracellular residence. In contrast, free HaloTag expressed from a plasmid did not exhibit increased immobilisation following phagocytosis of *E. coli*, confirming that the observed binding events reflect Pol1 activity rather than altered behaviour of the protein tag (Fig 1d). An additional advantage of HaloTag is that it can be labelled with ligands with distinct photophysical properties^21^. TMR fluorophores reversibly switch between bright and dark states, permitting recording of tens to hundreds of short-lived tracks (<1 s) per bacterium. Alternatively, the photostable fluorophore JFX650 provides fewer but long-lasting trajectories (>10 s) of individual molecules^23^. Using this approach, we observed single Pol1-Halo-JFX650 molecules diffusing within *E. coli* inside macrophages, becoming immobilised, and then dissociating again. The dwell times of individual binding events were consistent with previously measured kinetics of gap-filling DNA synthesis by Pol1 (e.g. 1.8 s in Fig. 1e), supporting the interpretation that binding events reflect active DNA repair^19^.

### Activity of BER and NER pathways increases in intracellular *E. coli*

Having established a method to visualise DNA repair activity in intracellular bacteria, we next investigated the contributions of specific repair pathways. NER removes helix-distorting DNA lesions, whereas BER employs lesion-specific DNA glycosylases to excise chemically modified bases^24^. Following lesion removal, both pathways generate gapped DNA intermediates that are filled by Pol1 ^3,8^. To assess NER activity, we deleted *uvrB*, which is essential for lesion verification in the pathway (Fig 2a). Using SMT, we found that the fraction of immobile Pol1-Halo-TMR molecules was significantly reduced in the Δ*uvrB* mutant compared to wild type (Fig 2b). Therefore, NER activity generates a substantial proportion of the gapped DNA substrates bound by Pol1. Notably, this effect was only observed after macrophage phagocytosis and not in unstressed bacteria prior to infection, demonstrating that NER is specifically engaged in response to DNA damage within the macrophage environment. To quantify the contribution of BER, we deleted the 8-oxoguanine DNA glycosylases MutM and MutY, which remove oxidised guanine lesions generated by ROS (Fig 2a). The Δ*mutM* Δ*mutY* strain also exhibited a marked reduction in bound Pol1-Halo molecules relative the wild type, showing that many excision repair events during intracellular residence target 8-oxoguanine lesions (Fig 2c).

**Figure 2.**
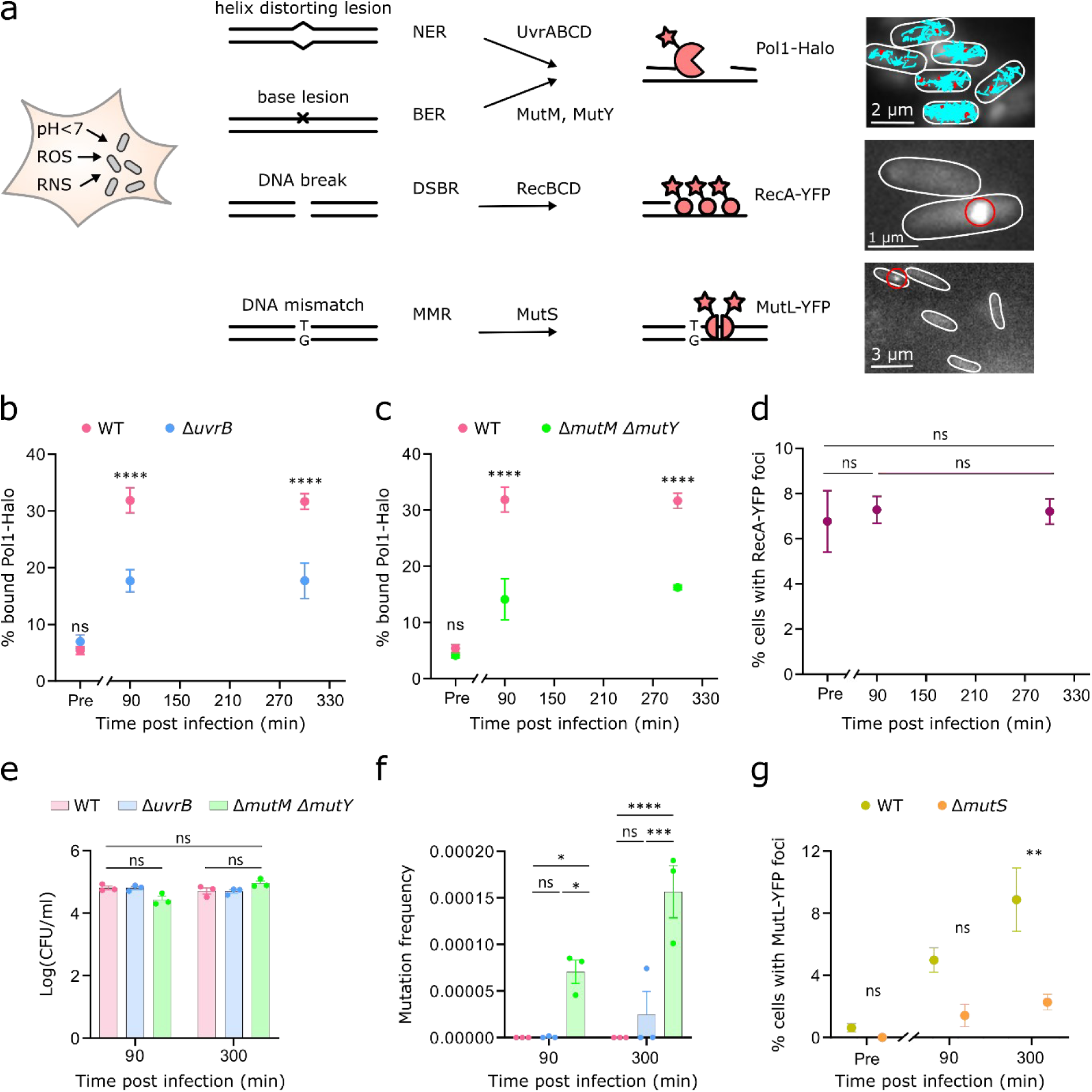
Macrophage phagocytosis triggers BER, NER, and MMR activity in *E. coli*. **(a)** Schematic overview of central DNA repair pathways in *E. coli*, with representative images of the corresponding reporter strains. **(b, c)** Percentages of immobile Pol1-Halo molecules pre-infection and at indicated times post-infection in the NER-deficient Δ*uvrB* isogenic mutant (b) and BER-deficient isogenic Δ*mutM* Δ*mutY* mutant (c) compared to wild type *E. coli*. **(d)** Percentages of cells with RecA-YFP foci pre-infection and at indicated times post-infection. **(e)** Quantification of intracellular bacterial survival, measured as colony-forming units (CFU), for wild type, BER-deficient, and NER-deficient strains recovered from lysed macrophages at the indicated times post-infection. **(f)** Mutation frequencies of wild type, BER-deficient, and NER-deficient strains from the proportion of rifampicin-resistant colonies relative to the total viable bacterial population recovered from lysed macrophages at the indicated times post-infection. **(g)** Percentages of wild type cells and MMR-deficient Δ*mutS* isogenic mutant cells with MutL-YFP foci pre-infection and at indicated times post-infection. Statistical analysis was performed using one-way ANOVA (d) or two-way ANOVA (b, c and e) followed by Tukey’s multiple-comparisons test. ns, not significant; \*\**P* < 0.01, \*\*\*\**P* < 0.0001. Error bars indicate mean ± S.E.M from 3 independent repeats.

To assess DSBR activity in intracellular *E. coli*, we utilised a RecA-YFP reporter that forms fluorescent foci at break sites^25^. RecA polymerisation also triggers autocleavage of the LexA repressor, thereby activating the SOS response and inducing a broader DNA damage regulon^26^. Despite the elevated BER and NER activities observed after phagocytosis, we detected no significant increase in RecA-YFP foci frequency in intracellular *E. coli* (Fig 2d). Furthermore, a strain carrying the non-cleavable *lexA3* allele, which constitutively represses the SOS response, showed intracellular survival indistinguishable from wild type *E. coli* (Extended Data Fig. 1)^26^. Together, these findings indicate that phagocytosis does not strongly induce DSBR or SOS activation in this model. This contrasts with pathogenic *E. coli* and *Salmonella* strains that are adapted to intracellular replication^27,28^. One potential explanation is that DSB formation depends on active DNA replication. As this strain of *E. coli* does not actively replicate within macrophages, base damage may be processed largely through excision repair without conversion into DSBs.

### Loss of DNA excision repair increases mutagenesis

Having established that BER and NER actively repair DNA damage incurred within the phagocyte environment, we next examined the functional importance of this activity. Deletion of *uvrB* or *mutM mutY* in *E. coli* did not detectably impair intracellular survival compared to wild type (Fig 2e). This is consistent with previous studies showing that disruption of BER or NER does not markedly sensitise *E. coli* to killing by ROS^14,29^. Thus, although excision repair pathways are engaged following phagocytosis, their loss does not substantially reduce *E. coli* survival under these conditions^30^. We therefore asked whether their principal function is to preserve genome integrity rather than immediate viability as the phagocyte environment is known to be mutagenic to bacteria^5^. We found that mutation frequencies of bacteria recovered from macrophages were elevated, particularly in the Δ*mutM* Δ*mutY* strain, relative to wild type (Fig 2f). This is consistent with the expectation that, in the absence of MutM and MutY, 8-oxoguanine lesions persist and mis pair with adenine during DNA synthesis^3^.

To directly quantify DNA synthesis errors in intracellular *E. coli*, we utilised a fluorescent MutL-YFP reporter. MutL is recruited to mismatches via the MMR pathway, forming foci that serve as a proxy for mutagenesis^31^. We observed a gradual increase in MutL-YFP foci frequency after macrophage phagocytosis, indicating progressive accumulation of DNA mismatches (Fig 2g). In contrast, MutL-YFP focus formation was strongly reduced in a Δ*mutS* strain, consistent with the requirement of MutS for mismatch recognition and MutL recruitment^3^. This shows that most MutL-YFP foci observed in wild type cells represent DNA mismatches. We conclude that intracellular residence within macrophages drives mutagenesis in *E. coli*, and that BER, NER and MMR pathways counteract DNA damage to preserve genome integrity.

### Macrophage internalisation slows protein diffusion in *E. coli*

While analysing Pol1 binding behaviour, we also observed a pronounced reduction in the mobility of diffusing Pol1 molecules following phagocytosis. The average diffusion coefficient of Pol1-Halo-TMR decreased by approximately twofold in intracellular *E. coli* (Fig 3a). Closer inspection revealed that protein motion within intracellular bacteria became spatially heterogeneous. Cumulative distribution frequencies and mean squared displacement (MSD) analysis showed sub-diffusive behaviour (Fig 3b, Fig 3c), and trajectories appeared clustered rather than uniformly distributed across the cytoplasm (Fig. 3d, e). The diffusion of free HaloTag also slowed in intracellular bacteria (Extended Data Fig. 2), suggesting that this effect reflects a general rigidification of the cytoplasm.

**Figure 3.**
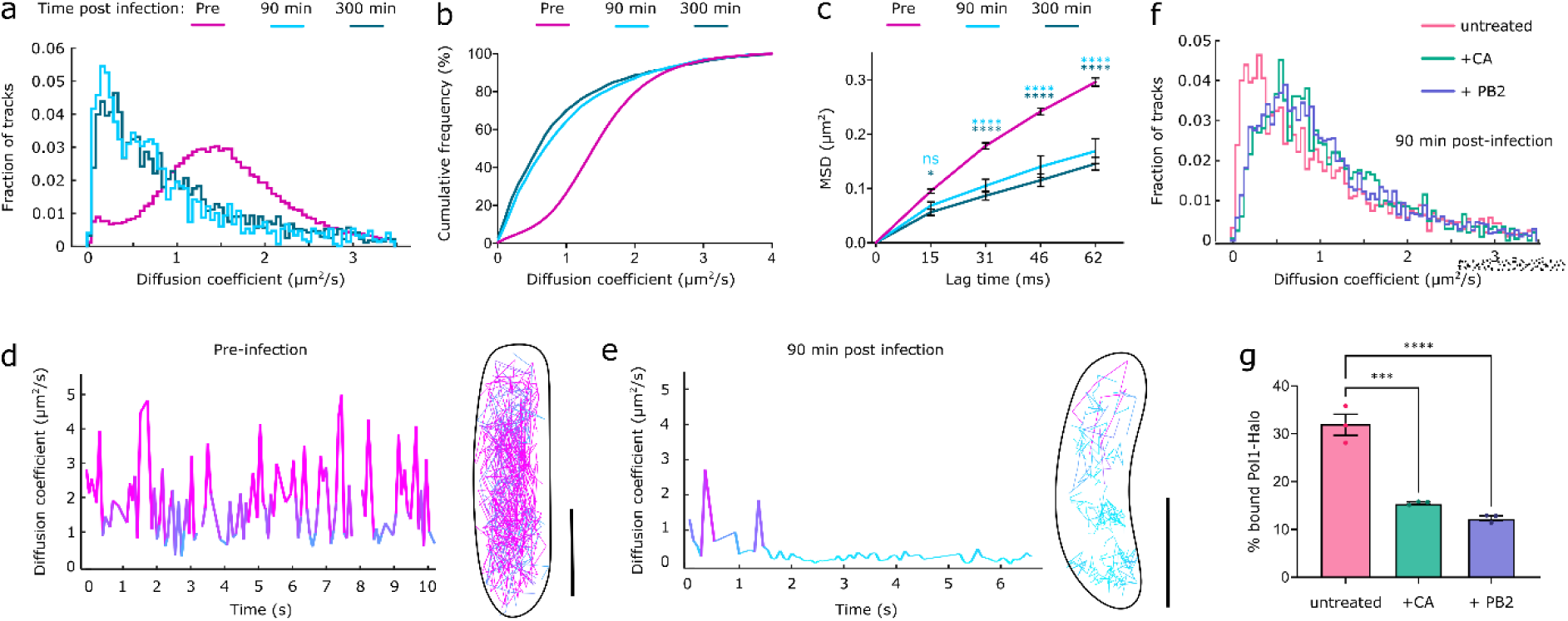
Macrophage internalisation reduces Pol1 mobility in intracellular *E. coli*. **(a,b)** Diffusion coefficient histograms and cumulative distributions of Pol1-Halo in *E. coli*, showing reduced mobility from pre-infection (pink) to 90 min (turquoise) and 300 min (teal) post-infection. Combined data from 3 independent repeats. **(c)** Mean squared displacement (MSD) curves of Pol1-Halo tracks showing increased motion confinement from pre-infection (pink) to 90 min (turquoise) and 300 min (teal) post-infection. (**d, e)** Representative time-traces of diffusion coefficients and corresponding tracks of single Pol1-Halo-JFX650 molecules in *E. coli* pre-infection and 90 min post-infection with segments of faster movement shown in pink gradient. Scale bars = 1 µm **(f)** Diffusion coefficient histograms of Pol1-Halo in *E. coli* at 90 min post-infection in macrophages treated with either concanamycin A (CA, green) or procyanidin B2 (PB2, blue) compared to untreated macrophages (pink). Combined data from 3 independent repeats. **(g)** Percentages of immobile Pol1-Halo molecules in *E. coli* at 90 min post-infection in macrophages treated with either concanamycin A (CA, green) or procyanidin B2 (PB2, purple) compared to untreated macrophages (pink). Statistical analysis was performed using two-way ANOVA followed by Tukey’s multiple-comparisons test (c) or one-way ANOVA followed by Dunnett’s multiple-comparisons test (f). ns, not significant; \**P* < 0.05, \*\**P* < 0.01, \*\*\**P* < 0.001, \*\*\*\**P* < 0.0001. Error bars indicate mean ± S.E.M. from 3 independent repeats.

A potential concern is that the reduced mobility of diffusing Pol1 could inflate the apparent immobile fraction, if slowly diffusing or transiently confined molecules were misclassified as DNA-bound. Several observations argue against this possibility and show that Pol1 binding, and the general slowdown are separable effects. Slowly diffusing Pol1 molecules remained clearly mobile, exploring small cytoplasmic regions and frequently transitioning between fast and slow movement (Fig 3e). Free HaloTag showed no increase in immobile fraction despite a slowdown after phagocytosis (Fig. 1d). Most directly, Δ*uvrB* and Δ*mutM* Δ*mutY* deletions reduced the fractions of immobile molecules (Fig. 2b, c) while the mobile molecules showed a similar slowdown as in wild type cells (Extended Data Fig. 2), demonstrating that the immobile population reports DNA repair activity rather than cytoplasmic state.

### Phagosome acidification and ROS production contribute to DNA damage and cytoplasmic rigidification

To identify host cell factors responsible for DNA damage and cytoplasmic rigidification, we treated macrophages with compounds that disrupt key antibacterial functions. Procyanidin B2 (PB2) reduces ROS production and shifts macrophages toward a less bactericidal, anti-inflammatory state^32,33^, whereas concanamycin A (CA) inhibits phagosome acidification by blocking the V-ATPase proton pump^34–36^. Treatment with either PB2 or CA partially restored Pol1 mobility and relaxed motion confinement (Fig 3f). Furthermore, PB2 and CA treatments lowered the fraction of immobile Pol1-Halo-TMR molecules, indicating that ROS production and phagosome acidification are sources of DNA damage in intracellular bacteria (Fig 3g). Mutation frequencies were also reduced in bacteria recovered from PB2-treated macrophages (Extended Data Fig. 3). Together, these results show that DNA damage and cytoplasmic rigidification are consequences of host cell defences, including oxidative and acid stress.

### Macrophage internalisation alters nucleoid organisation and cytoplasmic ultrastructure in intracellular *E. coli*

We considered whether changes in DNA density contribute to the observed diffusion slowdown, given that the mobility of DNA-binding proteins, such as Pol1, is influenced by frequent non-specific interactions with DNA^37^. Nucleoids labelled with a fluorescent HU-GFP fusion showed signs of hyper-compaction in intracellular bacteria (Extended Data Fig. 4), consistent with the nucleoid compaction observed in bacteria treated with DNA damaging agents *in vitro*. However, this effect was highly variable between bacteria in macrophages and became apparent only after 300 min post-phagocytosis, most prominently in bacteria showing signs of structural decay. Although nucleoid compaction may contribute to reduced mobility in some cells, its variability and delayed onset suggest that broader cytosolic changes underlie the general slowdown in protein diffusion.

We therefore turned to cryogenic electron tomography (cryo-ET) to examine intracellular bacteria and their host environment at ultrastructural resolution^38,39^. Macrophages grown on electron microscopy grids were infected with *E. coli* and plunge frozen. Cryogenic fluorescence imaging was used to identify intracellular bacteria and guide selective focused ion beam milling to generate ∼200 nm thin lamella. Cryo-ET revealed bacteria enclosed within phagosomes and surrounded by host lipid droplets, consistent with inflammatory activation of infected macrophages^40^ (Fig. 4). We compared extracellular bacteria grown on grids (Fig. 4a) with bacteria at 90 min and 300 min post-infection (Fig. 4b, c). In all samples, the bacterial outer and inner membranes were clearly resolved. Extracellular bacteria showed a smooth appearance with a relatively even distribution of macromolecules, reflecting the ribosome-rich cytoplasm (Fig. 4a). In contrast, intracellular bacteria showed progressive loss of homogeneity in the cytoplasm, with denser patches of material and regions of reduced density (Fig. 4b, c). At 300 min post-infection, we observed bacterial cell envelope damage and signs of nucleoid compaction as toroidal regions devoid of ribosome or other macromolecule density (Fig. 4c). These structural changes accompany the slowdown and confinement of protein diffusion observed by SMT, consistent with macrophage-induced stresses altering the physical properties of the bacterial cytoplasm.

**Figure 4.**
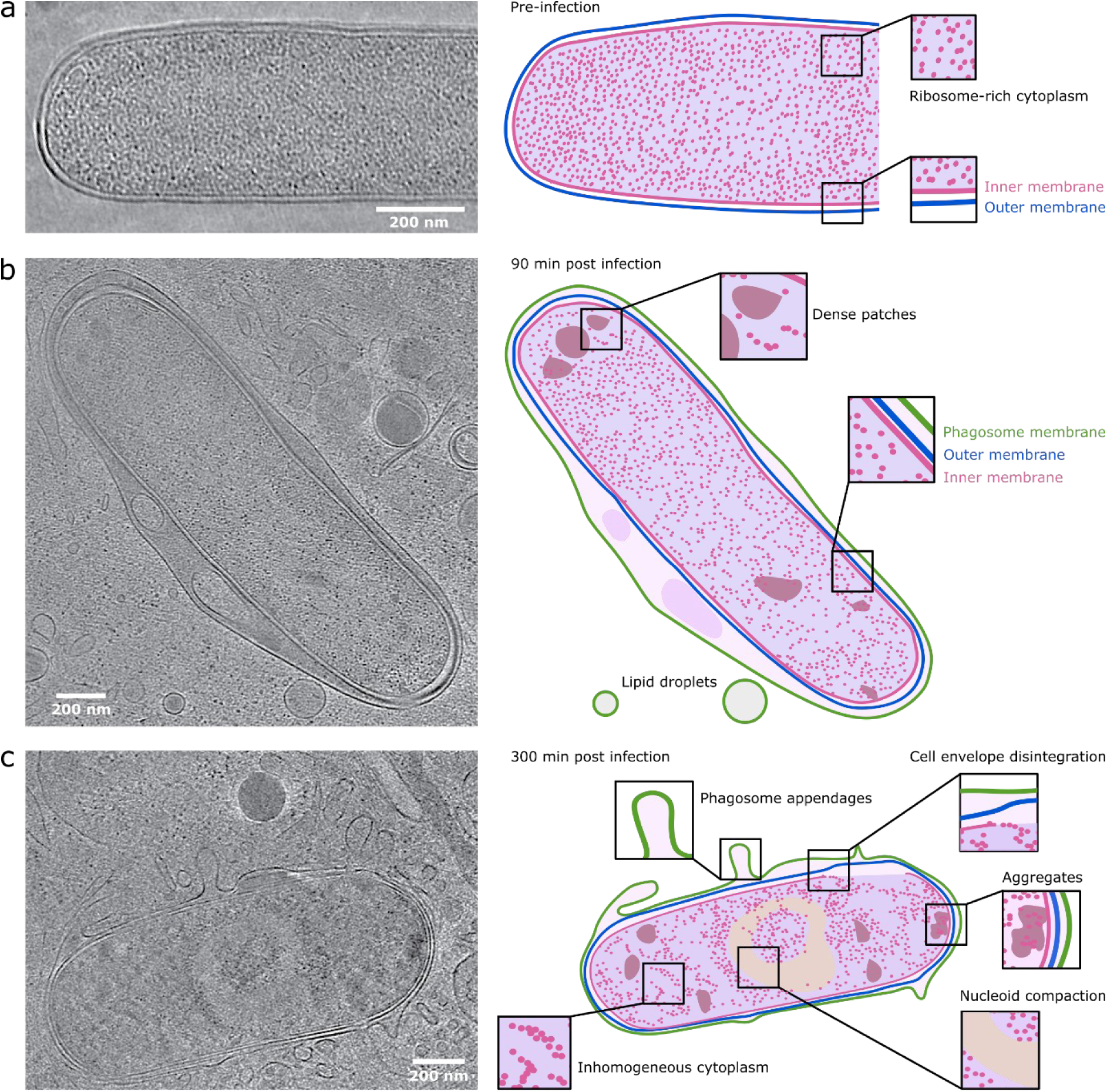
Macrophage internalisation alters cytoplasmic structure of *E. coli*. Representative cryo-ET slices of *E. coli* **(a)** pre-infection and at **(b)** 90 min and **(c)** 300 min post-infection. Corresponding schematics highlight structural features seen in tomograms.

### Cell-to-cell heterogeneity in cytoplasmic rigidification correlates with bacterial metabolic activity

Previous *in vitro* studies have shown that stress-induced transformations of the bacterial cytoplasm can profoundly affect cellular physiology^41,42^. Acidic pH, oxidative stress, and starvation alter macromolecular diffusion, protein aggregation, and nucleoid organisation, collectively shifting the cytoplasm toward a rigid, glass-like state^41^. This transition can protect cellular components, while reduced encounter rates between enzymes and substrates could slow metabolism and promote dormancy. Our results suggest that a similar cytoplasmic transition occurs in intracellular bacteria following phagocytosis, potentially representing an early physicochemical response that reduces metabolic activity.

We noticed cell-to-cell heterogeneity in the extent of nucleoid compaction and protein mobility changes, even among individual bacteria within the same macrophage. To assess whether this heterogeneity relates to metabolic state, we used the Min system as a genetically encoded, non-perturbative reporter compatible with intracellular measurements. In growing bacteria, MinCDE proteins dictate the site of cell division through ATP-dependent pole-to-pole oscillations^43^. Because oscillation dynamics depend on ATP availability, monitoring MinD-GFP by time-lapse imaging has been used as a proxy for cellular metabolic activity^44,45^. While ∼99% of bacteria pre-infection showed regular MinD-GFP oscillations, intracellular populations displayed a mixture of oscillating and non-oscillating bacteria (Fig 5a,b). The fraction of bacteria with MinD oscillations decreased over time within macrophages, indicative of progressive metabolic arrest under host-imposed stress (Fig 5c). In bacteria that retained oscillations, the period of pole-to-pole MinD-GFP movement increased significantly, consistent with reduced ATP availability and slower protein diffusion (Fig 5d). We then performed single-molecule tracking of Pol1-Halo-TMR after time-lapse imaging of MinD-GFP to directly link metabolic state and protein mobility in the same cells (Fig 5e). The slowdown in Pol1 diffusion was markedly more pronounced in bacteria lacking MinD-GFP oscillations, showing that cytoplasmic rigidification is closely associated with the loss of MinD oscillations (Fig 5f, g).

**Figure 5.**
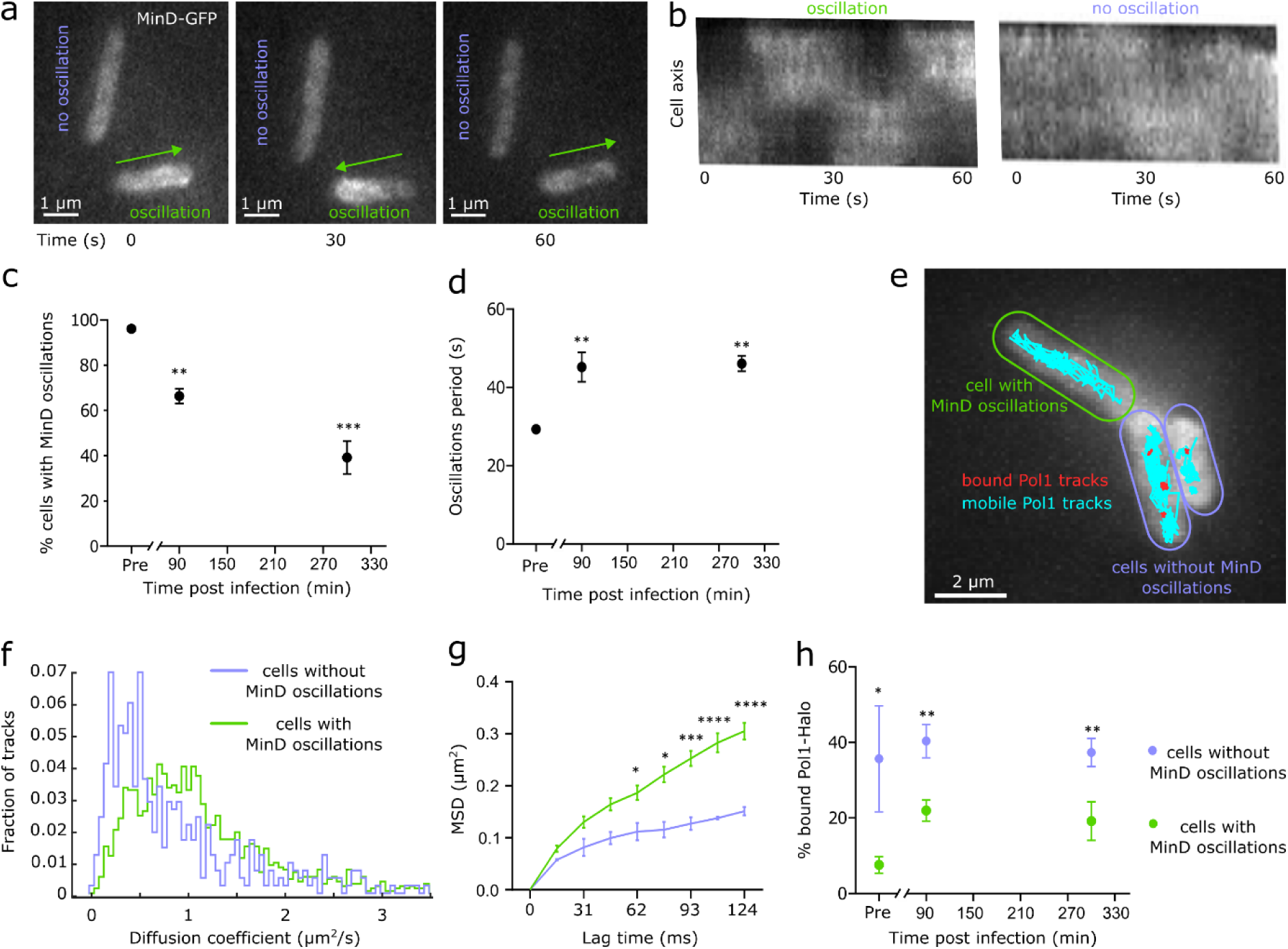
Coupling of MinD oscillation status, cytoplasmic rigidification and Pol1 binding in single intracellular *E. coli*. **(a)** Representative snapshots from a time-lapse movie showing *E. coli* cells with (green) and without MinD-GFP oscillations (purple) inside macrophage 90 min post-infection. **(b)** Kymographs showing MinD-GFP fluorescence along the long cell axis over time for *E. coli* cells with (green) and without oscillations (purple) inside macrophage 90 min post-infection. **(c)** Percentage of *E. coli* cells with MinD oscillations pre-infection and at indicated times post-infection. **(d)** Period of MinD oscillations from *E. coli* cells with MinD oscillations pre-infection and at indicated times post-infection. **(e)** Combined imaging of MinD-GFP and Pol1-Halo tracking in the same *E. coli* cells 300 min post-infection. Pol1-Halo tracks classified as mobile (turquoise) or immobile (red) show cell-to-cell heterogeneity in binding, related to presence (green) or absence (purple) of MinD oscillations. **(f)** Distributions of diffusion coefficients showing slowdown of Pol1-Halo diffusion in cells without (purple) compared to with (green) MinD oscillations 90 min post-infection. **(g)** Mean squared displacement (MSD) curves of Pol1-Halo showing increased motion confinement in cells without (purple) compared to with (green) MinD oscillations 90 min post-infection. **(h)** Percentages of bound Pol1-Halo molecules showing elevated binding in cells without (purple) compared to with (green) MinD oscillations pre-infection and at indicated times post-infection. Grouped data were analysed using two-way ANOVA followed by Tukey’s multiple-comparisons test. Comparisons of column data were performed using one-way ANOVA followed by Dunnett’s multiple-comparisons test against the 0 min post-infection control. \**P* < 0.05, \*\**P* <0.01, \*\*\*\**P* < 0.001, \*\*\*\**P* < 0.0001. Error bars indicate mean ± S.E.M. from 3 independent repeats.

### Elevated Pol1 binding in metabolically inactive cells

The observed heterogeneity in cytoplasmic state raised the question of whether DNA damage levels are similarly heterogeneous and related to the metabolic activity of individual bacteria. Single-cell analysis of Pol1 binding revealed variation in the fraction of bound molecules (Fig. 5e), indicating heterogeneity in repair protein engagement among intracellular bacteria. Combining MinD-GFP imaging with Pol1-Halo-TMR tracking in the same cells showed that this variability depends on metabolic state. Strikingly, bacteria lacking MinD oscillations exhibited substantially higher fractions of bound Pol1 molecules (Fig 5h). Cells that retained MinD oscillations also showed elevated DNA repair activity compared to bacteria pre-infection, but to a lesser extent than metabolically inactive cells (Fig 5h). While all intracellular bacteria engage in repair of macrophage-induced DNA damage, the subpopulation that has ceased MinD oscillations shows the highest fraction of bound Pol1. Notably, the small subpopulation of bacteria lacking MinD oscillations pre-infection (∼1%) also showed elevated Pol1 binding, and this subpopulation expands substantially within macrophages (Fig. 5d). Together, these findings reveal a connection between heterogeneous metabolic activity, cytoplasmic state and DNA repair protein engagement at the single-cell level.

## Discussion

Stress tolerance is a critical requirement for bacterial residence within a host, particularly following uptake by professional phagocytes^2^. Here, we focused on basal protective mechanisms provided by conserved DNA repair pathways. Using non-pathogenic *E. coli* as a model allowed us to investigate DNA repair functions in the absence of specialised immune evasion or virulence mechanisms. We developed a live-cell single-molecule tracking approach to directly measure protein activity in intracellular bacteria. This extends the microscopy toolbox to study intracellular bacteria, which includes single-molecule imaging of secreted proteins and single-cell reporters^22,46,47^. Direct visualisation of molecular mechanisms and bacterial behaviour *in situ* complements existing genetic and biochemical assays that rely on extraction of bacteria from host cells^16,22,46,48^. We anticipate that this generalisable method will be broadly applicable to interrogate the molecular processes that define the phenotypic states of individual bacteria during infection, including processes such as DNA replication, cell division, and gene regulation that have been extensively characterised using single-molecule tracking *in vitro^49^*. More broadly, intracellular bacteria can act as functional probes of the phagosome environment, because the damage they accumulate and the stress responses and repair pathways they engage reflect the types and intensity of stresses present within host cells. Focusing on DNA repair, we find that BER, NER, and MMR pathways are engaged following phagocytosis of *E. coli* by macrophages and that their loss increases bacterial mutagenesis. Exploiting the single-cell resolution of this approach further revealed that DNA repair activity varies substantially between individual bacteria. Notably, this heterogeneity is observed at the level of protein activity rather than the more commonly measured variation in gene expression. While phenotypic heterogeneity has previously been linked to differential expression of virulence factors and stress responses in intracellular pathogens, our results show that even in the absence of such specialised programmes, variability in genome maintenance is a general feature of bacteria exposed to host defences^4,17,18^.

As this heterogeneity was evident even between bacteria within the same macrophage, it is unlikely to be explained primarily by host cell state alone; instead, it points to differences in bacterial state and local microenvironment. By combining single-molecule tracking of Pol1 with MinD oscillations as a metabolic reporter, we show that heterogeneous DNA repair activity is coupled with metabolic state of individual bacteria, with inactive bacteria showing the highest fraction of bound Pol1. Starved bacteria may accumulate DNA damage if their ability to detoxify ROS and other host stressors is impaired. Additionally, reduced metabolic activity may constrain energy-dependent steps of DNA repair, potentially limiting repair efficiency. In parallel, we observe a pronounced reduction in protein mobility driven by macrophage-imposed stresses, including acidification and ROS. Cytoplasmic rigidification has been linked to pH-dependent changes in protein charge and macromolecular interactions, creating a glass-like state characterised by reduced mobility and increased molecular confinement^41,50^. Such cytoplasmic rigidification has been associated with reduced metabolic activity^41,51^, and may have broad consequences for intracellular processes that are driven by molecular encounters, including slowing the speed of lesion search by DNA repair proteins^37,52^. Together, our findings reveal how genome maintenance, cytoplasmic physical state, and metabolic activity are interconnected in bacteria exposed to phagocyte attack. Since elevated mutagenesis of bacteria within phagocytes could accelerate the emergence of drug-resistant and other adaptive variants, this heterogeneity may also have consequences for bacterial evolution during infection^53^.

## Methods

### *E. coli* genetic modification

All bacterial strains used in this study are derivatives of *E. coli* MG1655. Strains are shown in Supplementary Table 1 and all relevant PCR primers used for genetic modifications in Supplementary Table 2. Bacterial genome modifications were performed using Lambda Red recombination. All strains for SMT were constructed in a strain expressing a constitutive RecA-YFP chromosomal fusion, flanked by a FRT::kanR::FRT cassette. RecA-YFP was added next to the native locus using P1 transduction. The kanamycin resistance cassette was subsequently removed by expressing Flp recombinase from the temperature sensitive plasmid pCP20, transformed by heat shock. To create the Pol1-Halo fusion, the HaloTag sequence and a flexible 11-amino acid linker were added to the C-terminus of the endogenous *polA* gene using 50 nts homology overhangs, followed by a kanamycin resistance cassette flanked by frt sites. The gene fusion was moved into strain *E. coli* MG1655 RecA-YFP using P1 phage transduction. Free HaloTag was cloned into plasmid pBAD24 using isothermal assembly and transformed into MG1655 by heat shock. Chromosomal fluorescent protein fusions (MinD-sGFP, HU-GFP, MutL-mYFP, Tn7::mKate2) and gene deletions or mutations (Δ*uvrB*, Δ*mutM*, Δ*mutY*, *lexA3*), were moved to their respective backgrounds using P1 transduction (Supplementary Table 1).

### Cell culture and macrophage differentiation

THP-1 monocytes were propagated in Roswell Park Memorial Institute (RPMI) medium containing 10% (v/v) heat inactivated foetal bovine serum (FBS), and 1% (v/v) GlutaMAX. Monocytes were differentiated into macrophages using 200 nM phorbol 12-myristate-13-acetate (PMA) for 48h at 37°C, 5% CO2. Cells were rested for 24h at 37°C, 5% CO_2_ in RPMI medium without PMA prior to *E. coli* infection. Procyanidin B2 and Concanamycin A (Merck) were used at working concentrations of 10 µM and 1 µM, respectively, to pre-treat macrophages for 6h prior to infection and remained present in the culture medium throughout the subsequent infection period.

### *E. coli* infection of macrophages

*E. coli* cultures were grown in LB broth for 4-5h followed by M9 minimal media overnight. Minimal media was prepared by mixing M9 salts (15 g/l KH2PO4, 64 g/l Na2HPO4, 2.5 g/l NaCl, and 5.0 g/l NH4Cl), 2 mM MgSO4, 0.1 mM CaCl2, 0.5 mg/ml thiamine, MEM amino acids, 0.1 mg/ml L-proline, and 0.2% (v/v) glucose or glycerol. M9 cultures were generally grown with glucose, except for pBAD24::HaloTag harbouring strains which were grown with glycerol. Overnight cultures were diluted 1 in 100 and grown to an OD_600_=0.6, centrifuged, then resuspended in RPMI containing 10% (v/v) (FBS), and 1% (v/v) GlutaMAX. Macrophages were infected at a multiplicity of infection (MOI)=10 for 1h at 37°C, 5% CO_2_, followed by the addition of 200 µg/ml gentamicin for 30 min. Cells were subsequently left with 20 µg/ml of gentamicin and were incubated further at 37°C, 5% CO_2_ until designated time points for imaging or lysis. To calculate the number of viable bacteria, cells were lysed with 0.1% (v/v) Triton-X100 in phosphate buffered saline (PBS) for 5min at room temperature. Viable bacteria were enumerated by plating onto LB agar and counting colonies the next day. To quantify mutation frequency, bacteria recovered from macrophage lysates were cultured in liquid LB medium for 16h to expand the number of cells, then plated on LB agar with 100 μg/ml rifampicin and a 10^7^-fold dilution on LB agar without antibiotic. Mutation frequency was calculated as the proportion of rifampicin-resistant colonies relative to the total viable bacterial population.

### SMT in intracellular *E. coli*

1 x 10^6^ THP-1 macrophages were seeded onto MatTek 1.5 mm cover-glass tissue culture dishes and differentiated with PMA as described above. Prior to seeding, fluorescent background particles were removed from dishes using air plasma (Plasma Etch) and incubated with 1% (w/v) bovine serum albumin (BSA) for 10 min at 37°C/5% CO_2_.

*E. coli* MG1655 expressing Pol1-Halo (or free HaloTag) were streaked from frozen glycerol stocks onto LB agar containing the appropriate antibiotic. Bacteria further expressed RecA-YFP to allow identification within macrophages prior to SMT. For MinD imaging, cells alternatively expressed MinD-sGFP instead of RecA-YFP. A single colony was inoculated into LB broth containing the appropriate antibiotic and grown for 4h at 37°C, 200 rpm to OD_600_=∼0.8. Cultures were diluted 1 in 1000 in M9 minimal medium and grown overnight to stationary phase at 37°C, 200 rpm.

The next day, cultures were diluted 1 in 50 (v/v) in M9 minimal media and grown for 2h at 37°C. Cultures were centrifuged at 3000 *x g* for 10 min to form a pellet. The pellet was resuspended in 100 µl M9 minimal media containing 5 µl of 2.5 µM TMR ligand (Tetramethylrhodamine, for Pol1-Halo tracking, Promega), or 5 µl of 2.5 µM of JFX554 (showing superior brightness for free HaloTag tracking, Promega), or 0.5 µl of 50 µM JFX650 (for prolonged Pol1-Halo tracking, Janelia Farm) and incubated at room temperature for 30 min. Residual dye was removed by washing labelled cells thrice by centrifugation at 6000 *x g* for 3 min and resuspension in 1 ml M9 minimal medium. The culture was allowed to recover for 30 min at 37°C, 200 rpm. Bacteria were subsequently enumerated by measuring OD_600_, resuspended in complete RPMI, and added to adherent macrophages in plasma-cleaned cover-glass tissue culture dishes at a MOI of 10. Infected macrophages were incubated at 37°C/5% CO₂, with the start of infection designated as 0h. Following 1h of incubation, extracellular bacteria were removed, and the macrophages were washed twice with complete RPMI containing 200 µg/mL gentamicin. The macrophages were then incubated in complete RPMI containing 200 µg/mL gentamicin for 30 min at 37°C/5% CO₂. At 90 min post-infection, macrophages were washed again and incubated until the defined experimental time points in complete RPMI containing 20 µg/mL gentamicin at 37°C with 5% CO₂. Following removal from the tissue culture incubator, culture dishes were imaged for 1h. The reported time points of 90- and 300-min post-infection correspond to the beginning of each 1h imaging session.

Pre-infection bacteria were grown and labelled as described above but, rather than being used for macrophage infection, were imaged on agarose gel pads. The pads were prepared by combining Molecular Biology Agarose (Bio-Rad) with M9 minimal medium at a final agarose concentration of 1% (w/v) and warmed in a microwave until the agarose had dissolved. Around 1 ml of the resulting molten M9 agarose was transferred onto a #1.5 borosilicate cover glass (VWR International). A second plasma-cleaned cover glass was placed over the agarose to form a uniform pad, which was then allowed to set. Just before microscopy, the upper cover glass was lifted off, and the cells were applied directly to the surface of the solidified agarose. The preparation was subsequently covered with a fresh plasma-cleaned cover glass for SMT data acquisition. For Procyanidin B2 or Concanamycin A treatments of bacteria pre-infection, bacteria were incubated for 1h at 37°C with the compounds at the same concentrations as for macrophage treatments before SMT data acquisition on agarose pads.

### SMT data acquisition

SMT experiments were conducted on a custom-built inverted optical microscope with ASI (Applied Scientific Instrumentation) and Thorlabs components configured for total internal reflection fluorescence (TIRF) imaging. Fluorescence excitation was provided by a multi-laser engine (iChrome MLE, Toptica Photonics) capable of delivering wavelengths at 405, 488, 561, and 640 nm. Excitation and emission paths were separated using a quad-band dichroic mirror (ZT405/488/561/640rpc, Chroma). The microscope was equipped with a Nikon Plan Apo λ 100×/1.45 NA oil immersion objective. Fluorescence emission for each channel was selected via a motorized filter wheel containing: 425–475 nm and 500–550 nm bandpass filters (Chroma) for 405 nm and 488 nm excitation respectively, 575 nm long-pass filter (Chroma) for 561 nm excitation and 655 nm long-pass filter (Chroma) for 640 nm excitation.

The detection system consisted of an EMCCD camera (iXon Ultra 897, Andor) controlled via Solis software, with 300x EM gain and an effective pixel size of 107 nm. Movies were captured with an exposure time of 15 ms/frame (Pol1-Halo) or 10 ms/frame (free HaloTag). The additional camera readout time of 0.475 ms gave frame intervals of 15.48 ms or 10.48 ms respectively.

A TG-1000 controller (ASI) interfaced with multiple motorized ASI components including the XY translation stage for sample positioning, Z and piezo stages for axial adjustments, a dichroic mirror slider, emission filter wheel, and an LED condenser. All microscope components apart from the camera were operated using Micro-Manager software. For background suppression, the system incorporated a TIRF illumination module based on a translatable lens stage (MB1530F/M, Thorlabs). This stage, driven by a motorized actuator (Z812, Thorlabs) and Kinesis KDC101 controller, allowed precise adjustment of the laser beam position at the objective back focal plane to achieve variable-angle illumination. An Okolabs chamber maintained the sample at a constant temperature of 37°C during imaging.

Regions of interest containing a suitable density of bacteria were selected by recording snapshots of RecA-YFP fluorescence under 0.01 Wcm^2^ 488 nm laser illumination. Alternatively, for experiments involving MinD-GFP, movies were recorded under 0.01 Wcm^2^ 488 nm laser illumination with 10 ms exposure time. Next, a transmitted light image of macrophages was captured under LED illumination. Subsequently, Halo-TMR (or Halo-JFX554) was excited with 2.6 Wcm^2^ 561 nm laser illumination or Halo-JFX650 with 640 nm laser illumination. Movies of 5,000–10,000 frames were recorded after briefly deactivating fluorophores such that the density of fluorescent spots reduced to less than one spot per cell per frame.

### SMT Analysis

Microscopy data were analysed in MATLAB R2025b. Outlines of bacteria were segmented based on the RecA-YFP fluorescent snapshot using MicrobeTracker, with only cells displaying both poles within the imaging plane being chosen for downstream analysis. Fluorescent spots were localised in each frame of SMT movies using a phasor localisation detection algorithm, using a minimum intensity threshold to discern fluorescent signals from background. Detected spots were linked together to form single-molecule trajectories provided each molecule within adjacent frames were within a radius of 8 pixels from the prior frame, and they originated from the same cell, as defined by segmentation analysis. Each trajectory was assigned a one-frame memory, allowing localizations separated by a single undetected frame to remain linked if all other criteria were met. For trajectories lasting at least 5 frames, the first 4 frame-to-frame steps were used to calculate the mean squared displacement (MSD). The MSD alongside the time between consecutive frames (*Δt*) was used to calculate the diffusion coefficient for each trajectory:

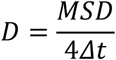

Based on previous investigations of Pol1 diffusion behaviour^19^, we estimated the proportions of molecules in bound and diffusing states, by fitting the distribution of *D* values to a sum of two probability density functions,:

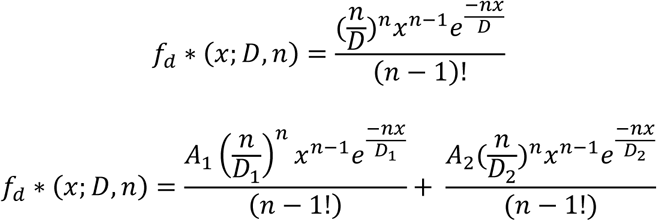

Here, f is the probability of measuring a molecule with diffusion coefficient x; D_1_ and D_2_ are the average diffusion coefficient of each state; n is the number of steps the MSD is calculated over; and the proportions of molecules in each diffusive state satisfy A_1_+A_2_= 1. The diffusion coefficient of the bound population was fixed at D_1_ = 0.17 µm^2^/s, so the fit estimated the values of the free parameters D_2_ and A_1_.

### EPI fluorescence microscopy and analysis

Nucleoid imaging (HU-GFP), Tn7::mKate2, and mismatch repair foci quantification (MutL-YFP) was performed using EPI fluorescence microscopy. Macrophages were infected, as described above, and fixed at the stated time points with 4% (v/v) formaldehyde solution in PBS. Snapshot images were captured on a Nikon Ti Eclipse Inverted fluorescent microscope with Photometrics Prime 95B camera and Lumencor LED illumination. Exposure times were 500 ms for MutL-YFP (λ=508 nm) and 1s for HU-GFP (λ=470 nm) using 50% of maximal LED excitation intensities.

### Cryogenic electron tomography

1x10^6^ monocytes were seeded onto EM grids (R2/1 Au 200 mesh, Quantifoil) coated with 1 µg/ml human fibronectin. Cells were differentiated with PMA and infected as described above. Bacteria were grown in the presence of 1 µg/ml MitoTracker Green FM dye to stain the bacterial inner membrane for cryogenic fluorescence imaging. At the chosen time after infection, 5 µg/ml Hoechst 33342 was added to the cells and incubated for 5 min at RT. Following incubation, cells were washed thrice in PBS, and grids were back-blotted for 5 s and then plunge-frozen in a liquid ethane-propane mixture. Intracellular bacteria were assigned approximate grid coordinates by cryogenic fluorescence imaging using a Leica STELLARIS 5 cryo-confocal microscope. Bacterial cells near the stained nucleus were selected as likely intracellular candidates. Cryo-FIB lamellae were prepared using a Crossbeam 350 dual-beam cryo-FIB scanning electron microscope (cryo-FIB-SEM, Zeiss) equipped with a Leica cryo-transfer system and cryo-stage. Lamellae were milled to 500 nm with a reducing current (7 nA – 300 pA) and then polished to ∼200 nm at 50 pA.

All cryogenic electron tomography (cryo-ET) datasets were collected on a Titan Krios transmission electron microscope (FEI, Thermo Fisher) operating at 300 kV and equipped with a Bio-Quantum K3 direct electron detector and energy filter (Gatan). Data acquisition was performed using SerialEM^54^. Tilt series were acquired with a bidirectional tilt scheme ranging from -60° to +60° starting from 0° with 3° increments at the nominal magnification of 6,500× corresponding to a pixel size of 1.312 nm. Tomograms were reconstructed using IMOD at bin 4, yielding a final pixel size of 5.248 nm^55^. Tomograms were de-striped and filtered for display using Leonardo Toolset^56^.

### Statistical analysis

Statistical analysis was performed using either MATLAB R2022b or GraphPad 10.5.0 and referenced in the corresponding figure legends. For significance testing, a threshold of α=0.05 was used. Normality (Gaussian distribution) was assessed by the Shapiro-Wilk test. If data was normally distributed then either a Student’s t-test, or the appropriate ANOVA was performed (unless otherwise stated) with multiple comparisons where appropriate and applying Geisser-Greenhouse correction if sphericity assumption was not met. For non-normally distributed data, which could not be transformed, non-parametric Kruskal-Wallis test analysis was performed.

## Data availability

Source data and MATLAB data analysis scripts associated with this data will be deposited in the Oxford University Research Archive (ORA, https://ora.ox.ac.uk/).

## Code availability

MicrobeTracker suite is an open-source software package for cell segmentation in MATLAB. The MATLAB script for diffusion analysis is included in the supplementary material.

## Acknowledgements

We thank all members of the Uphoff lab, past and present, alongside David Sherratt and Christoph Tang, for discussions. We thank Achillefs Kapanidis and Cees Dekker for providing bacterial strains. This research was funded by a Wellcome Trust Sir Henry Dale Fellowship (206159/Z/17/Z) and a Research Prize Fellowship by the Lister Institute of Preventative Medicine to S.U., and an Academy of Medical Sciences Springboard Award SBF008\1157 and a fellowship from the Kavli Institute for Nanoscience Discovery Oxford to L.B. F.A.S. acknowledges a Junior Research Fellowship at New College, Oxford. C.Q. acknowledges a Summer Studentship by the Lister Institute of Preventative Medicine.

## Author contributions

F.A.S., L.B. and S.U. conceived the project and interpreted the data. F.A.S. and S.U. performed experiments, supervised students, and analysed and curated the data. N.C.D. performed cryo-FIB/cryo-ET experiments alongside image processing. N.C.D., T.B.O’H. and L.B. analysed the cryo-ET data. F.A.S., C.K., C.Q. and S.U. generated mutants and performed the microscopy. E.H. and A.K. assisted with tissue culture and contributed to development of infection and imaging protocols. A.M. supported SMT measurements, instrumentation and analysis. F.A.S., N.C.D., L.B. and S.U. wrote the manuscript with feedback from all authors.

