## Supplementary information for "Single-molecule imaging of DNA repair and cytoplasmic rigidification in intracellular bacteria"

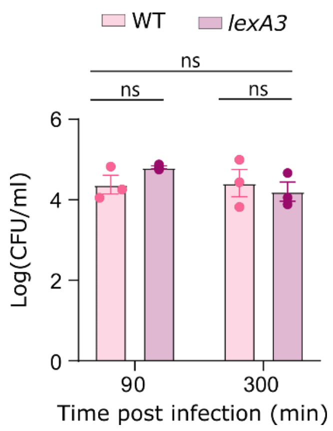


**Extended Data Figure 1. Survival of *E. coli* in THP-1 derived macrophages does not require SOS response induction**. Quantification of intracellular bacterial survival, measured as colony-forming units (CFU), for wild-type and a non-cleavable *lexA* derivative recovered from lysed macrophages at the indicated times post-infection. Statistical analysis was performed using two-way ANOVA followed by Tukeys multiple-comparisons. ns, not significant; Error bars indicate mean ± S.E.M from 3 independent repeats.

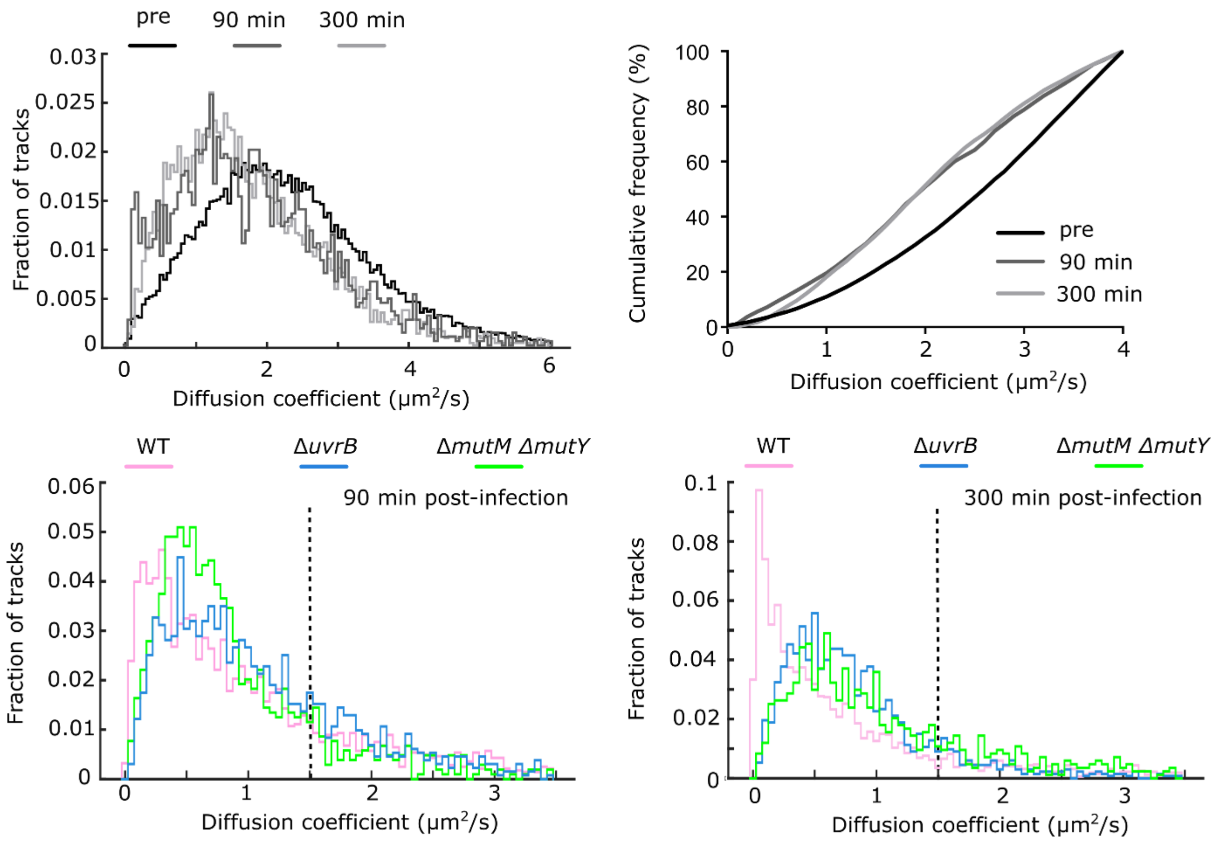


**Extended Data Figure 2. The macrophage environment induces a general slowdown of protein diffusion independently of changes in Pol1 binding. (a, b)** Diffusion coefficient histograms and cumulative distributions of free HaloTag in *E. coli*, showing reduced mobility from pre-infection (black) to 90 min and 300 min (grey scales) post-infection. Combined data from 3 independent repeats. **(c,d)** Distributions of diffusion coefficients showing slowdown of Pol1-Halo diffusion in wild-type (pink), NER-deficient (blue), and BER-deficient (green). The black dashed line represents the mean diffusion coefficient of Pol1-Halo pre-infection. Combined data from 3 independent repeats. The mobile Pol1 population undergoes a similar diffusion slowdown in wild-type and repair-deficient strains, whereas the immobile fraction is specifically reduced in the mutant strains. These results indicate that the infection-associated reduction in protein mobility occurs independently of Pol1 binding.

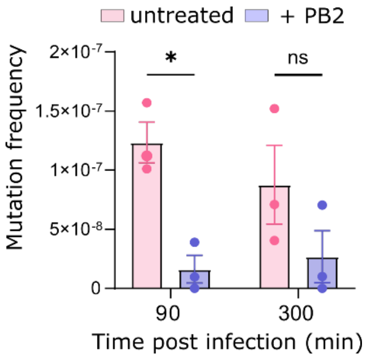


**Extended Data Figure 3. Reduced mutation frequencies in bacteria recovered from PB2-treated macrophages**. Mutation frequencies of *E. coli* recovered from untreated and PB2-treated macrophages. Mutation frequencies were calculated from the proportion of rifampicin-resistant colonies relative to the total viable bacterial population recovered from lysed macrophages at the indicated times post-infection. Statistical analysis was performed using two-way ANOVA followed by Šídák's multiple comparisons. ns, not significant; **P* < 0.05. Error bars indicate mean ± S.E.M from 3 independent repeats.

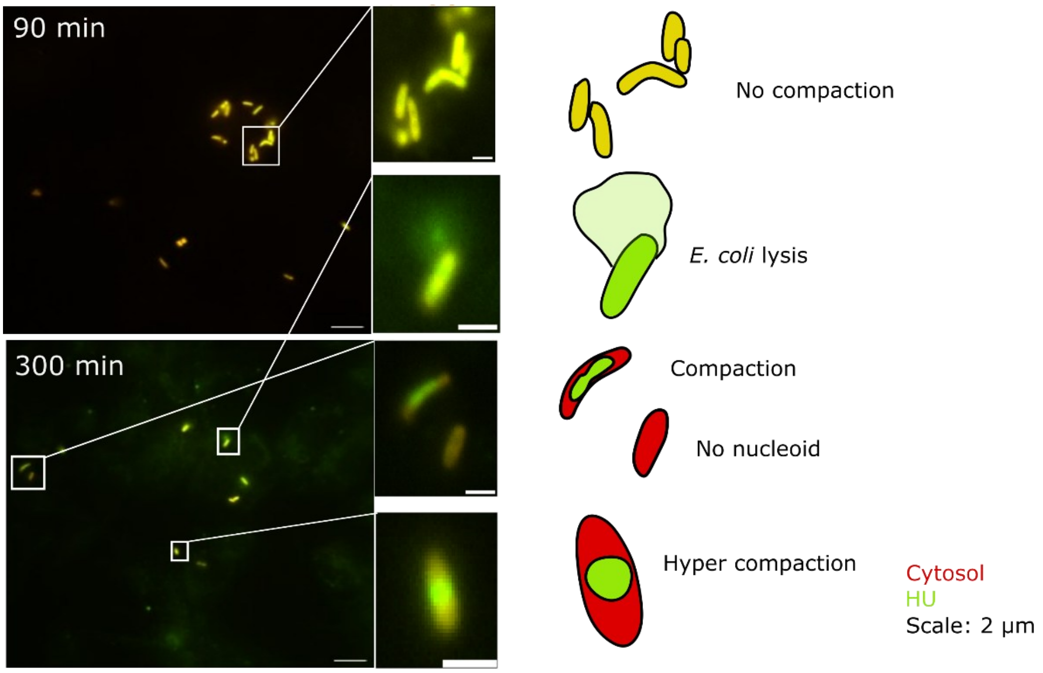


**Extended Data Figure 4. Nucleoids labelled with a fluorescent HU-GFP fusion show bacterial lysis and signs of hyper-compaction in intracellular *E. coli*.** Representative fluorescent images shown as overlay of HU-GFP (green) and Tn7::mKate2 (red) as a cytoplasmic marker of *E. coli* at (a) 90 min and (b) 300 min post-infection of macrophages. Corresponding schematics highlight observed structural features.

**Supplementary Table 1: Full list of strains used in this study**

| Perturbated pathway | Full Name of strain | Strain ID (as referred to in text) | Source |
|---|---|---|---|
| Wild type | *Escherichia coli* K-12 substr. MG1655 | MG1655 wild type | |
| | *E. coli* MG1655 RecA-YFP Pol1-Halo | MG1655 Pol1-Halo | Johan Elf lab[24]<br>Stephan Uphoff lab[20] |
| | *E. coli* MG1655 RecA-YFP pBAD24::Halo | MG1655 pBAD24::Halo | This study |
| | *E. coli* MG1655 MinD-GFP Pol1-Halo | MG1655 MinD-GFP Pol1-Halo | Cees Dekker lab[45] |
| | *E. coli* MG1655 MutL-mYPet mKate2 | MG1655 MutL-YFP | This study |
| | *E. coli* MG1655 HU-GFP mKate2 | E. coli MG1655 HU-GFP | Achillefs Kapanidis lab |
| NER | *E. coli* MG1655 Δ*uvrB* RecA-YFP Pol1-Halo | MG1655 Δ*uvrB* Pol1-Halo | CGSC* 8819 |
| BER | *E. coli* MG1655 Δ*mutM* Δ*mutY* RecA-YFP Pol1-Halo | MG1655 Δ*mutM* Δ*mutY* Pol1-Halo | CGSC* 11684<br>CGSC* 11685 |
| MMR | *E. coli* MG1655 Δ*mutS* MutL-mYPet mKate2 | MG1655 Δ*mutS* MutL-YFP | CGSC* 10126 |
| DSBR | *E. coli* MG1655 LexA3 | MG1655 *lexAG85D* | CGSC* K996 |

**CGSC: Coli Genetics Stock Centre*

**Supplementary Table 2: Primers used in this study**

| Construct | Primer ID |
|---|---|
| Pol1-Halo | Pol1_pSU005_F: TGCCGTTGCTGGTGGAAGTGGGGAGTGGCGAAAACTG GGATCAGGCGCACTCGGCTGGCTCCGCTGC<br>Pol1_pSU005_R: ACGTGACAGCTTATGTTGCTTACTTACGAAAAAAGGCAT GTTCAGGCGAATCTATGAATATCCTCCTTAG |
| pBAD24::HaloTag | pBAD24_backbone_F: GTACCCGGGGATCCTCTAG<br>pBAD24_backbone_R: CATGGTGAATTCCTCCTGC<br>pSU005_HaloTag_F: AGCAGGAGGAATTCACCATGGGATCCGAAATCGGT ACTG<br>pSU005_HaloTag_R: TCTAGAGGATCCCCGGGTACTTAACCGGAAATCTCCAG |
| Δ*uvrB* | pK3_Δ*uvrB*_F: AACTCCTTCAGGTAGCGACTGTGTAGGCTGGAGCT<br>pK3_Δ*uvrB*_R: CAGGATAGCGAAGAAGACTGAGGACCATGGCTAATT |
| Δ*mutM* | pK4_Δ*mutM*_F: CGACTAAACATGCGCAGCGGGCAACGTTTTATTGTCGGC AGTGCCAGAAGTCGGCTGGCTCCGCTGC<br>pK4_Δ*mutM*_R: CGACTAAACATGCGCAGCGGGCAACGTTTTATTGTCGGC AGTGCCAGAAGTCGGCTGGCTCCGCTGC |
| Δ*mutY* | pK4_Δ*mutY*_F: CGGCTCCCGTGGAGCGTTTGTTACAGCAGTTACGCACTG GCGCGCCGGTTTCGGCTGGCTCCGCTGC<br>pK4_Δ*mutY*_R: GTACAAAAAATCGTTCTGCTCATAAATCATCCTCTTTATCGA CTCACGCGTATGAATATCCTCCTTAG |
| Δ*mutS* | pK4_Δ*mutS*_F: TCACCCCGCGTCAGGCGCTGGAGTGGATTTATCGCTTGA AGAGCCTGGTGTCGGCTGGCTCCGCTGC<br>pK4_Δ*mutS*_R: GTCAGTTGTCGTTAATATTCCCGATAGCAAAAGACTATCGG GAATTGTTATATGAATATCCTCCTTAG |

**Summary of source data**

**Source Data Table 1:** SMT tracking metadata. Each condition was tested over 3 experimental replicates, consisting of 10-20 fields of view per replicate.

| *E. coli* strain or condition | Time Point | Average Bound Fraction (%) | Total Number of Tracks | Total Number of bacterial cells | Average Number of Tracks/bacterial cell |
|---|---|---|---|---|---|
| Pol1-Halo | Pre-infection | 5.5 | 46767 | 2238 | 20.90 |
| | 90 min | 31.8 | 1680 | 532 | 3.16 |
| | 300 min | 31.6 | 1006 | 333 | 3.02 |
| free-Halo | Pre-infection | 3.7 | 26928 | 1094 | 24.61 |
| | 90 min | 5.4 | 2893 | 102 | 28.36 |
| | 300 min | 6.7 | 1829 | 53 | 34.51 |
| Δ*uvrB* Pol1-Halo | Pre-infection | 9.9 | 9871 | 822 | 12.01 |
| | 90 min | 17.8 | 1163 | 95 | 12.24 |
| | 300 min | 18.5 | 626 | 118 | 5.31 |
| Δ*mutM* Δ*mutY* Pol1-Halo | Pre-infection | 4.1 | 160614 | 1925 | 83.44 |
| | 90 min | 14.1 | 1315 | 102 | 12.89 |
| | 300 min | 16.2 | 836 | 72 | 11.61 |
| Pol1-Halo +CA | Pre-infection | 6.06 | 93385 | 1455 | 64.18 |
| | 90 min | 18.1 | 6464 | 87 | 74.30 |
| Pol1-Halo +PB2 | Pre-infection | 6.2 | 33309 | 1597 | 20.86 |
| | 90 min | 12.4 | 1996 | 169 | 11.81 |

**Source data 2:** Quantification of mean diffusion coefficients (D) of Pol1-Halo.

| Condition | Variables | Mean | SEM | Lower 95% CI of Mean | Upper 95% CI of Mean | p-value |
|---|---|---|---|---|---|---|
| Pol1-Halo | Pre-infection | 1.510 | 0.003 | 1.503 | 1.517 | - |
| | 90 min | 0.987 | 0.016 | 0.955 | 1.018 | <0.0001 |
| | 300 min | 0.910 | 0.023 | 0.866 | 0.956 | <0.0001 |
| free Halo | Pre-infection | 2.499 | 0.008 | 2.485 | 2.515 | - |
| | 90 min | 2.063 | 0.016 | 2.030 | 2.096 | <0.0001 |
| | 300 min | 2.060 | 0.024 | 2.013 | 2.08 | <0.0001 |
| 90 min | Untreated | 0.987 | 0.016 | 0.955 | 1.019 | - |
| | +CA | 1.103 | 0.011 | 1.082 | 1.124 | <0.0001 |
| | +PB2 | 1.104 | 0.017 | 1.071 | 1.138 | <0.0001 |
| 90 min | MinD oscillating | 1.110 | 0.019 | 1.072 | 1.148 | - |
| | MinD non-oscillating | 0.870 | 0.036 | 0.798 | 0.942 | <0.0001 |
| 300 min | MinD oscillating | 1.114 | 0018 | 1.079 | 1.150 | - |
| | MinD non-oscillating | 0.909 | 0.031 | 0.848 | 0.971 | <0.0001 |
